# Electron Microscopic Identification of Distant Metastatic Cell in Human Prostate Cancer in Tissue section

**DOI:** 10.1101/787853

**Authors:** Akhouri A. Sinha

## Abstract

Mutation in chromatin/DNA/genes of stem cells is a prerequisite for the development of cancer in the benign prostate. Mutation imparts chronic proliferation advantage to invasive and metastatic cancer cells, but not in benign prostate. We hypothesize that identification of metastatic prostatic cancer (PC) cells in tissue sections can lead to an early diagnosis and treatment of patients, hopefully, reducing death due to this disease. We have tested our hypothesis using biopsy and/or radical prostatectomy specimens of untreated and DES (diethylstilbestrol)-treated prostate cancer and transmission electron microscopy. Materials and methods: Tissue samples embedded in Epon were thin sectioned and examined on RCA EMU 3 or 4 transmission electron microscope. Results: We have identified two types of PC stem cells. Invasive stem cell invades the adjacent stroma whereas metastatic cell, a lineage of stem cell, is dedifferentiated columnar/cuboidal cells. Metastatic cell is identified by nuclear plasticity (pleomorphic nucleus), loss of nuclear membranes, loss of boundary between nucleus and cytoplasm and presence of electron dense molecules. Nuclear chromatin/DNA appears as electron dense molecules which readily pass plasma and basement membranes and enter the capillary before adhering to the red cell surface. Electron dense molecules are found in capillary, undoubtedly, reach metastatic site (s). Discussion and conclusion: Chromatin/DNA are small molecules that can readily cross many barriers, unlike an individual cancer cell. We show that invasive a cell is stem cell and metastasis develops from dedifferentiated columnar/cuboidal nuclear electron dense molecules. This is the first study to cells characterize metastatic cell in tissue sections.

## Introduction

Mutation is required in DNA/genes of stem cells of benign prostate to develop prostate cancer (PC) and in benign organs of other solid organ cancers. Mutation imparts chronic proliferation advantage to invasive and metastatic cancer cells, but not in benign prostate (Ross et al., 2007). (Mucci et al., 2000). Many mutagens (such as pesticides, herbicides, toxins, chemicals, contaminated food and water) circulating in capillaries around prostate gland have the potential of inducing mutation in genes. Repeated exposures to mutagens produce deadly cancer. The number of mutated genes in PC is unknown in PC. Specific mutagens inducing PC are also unknown. Mutation varies in each type of cancer and produces heterogeneous disease (Hanahan and Weinberg, 2011 {Ramel, 1984 #1295). Mutation enables cancer cells to sustain chronic proliferation in invasive and metastatic cells whereas benign prostate and benign prostatic hyperplasia (BPH) do not have that advantage (Hanahan and Weinberg, 2011) (Bryden et al., 2001) (Scholzen and Gerdes, 2000) (Lindboe and Torp, 2002). Metastatic disease is responsible for majority of PC deaths (Van Etten and Dehm, 2016). Metastasis usually occurs in about 10% PC patients indicating metastatic inefficiency (Weiss, 1981).

A brief review of the vast literature on metastasis has shown that metastasis in humans is distinctly different from that shown in human and animal models and cell lines (Schirrmacher, 1985) (Fidler, 1989) (Heppner and Miller, 1989) (Liotta and Stetler-Stevenson, 1991). Metastasis occurs in nearly every solid organ cancer. Metastases vary greatly in human cancer patients, e.g. prostate, breast, colorectal, glioblastoma, and pancreatic cancers (Werb, 1989) (Fidler, 1989) (Tarin, 1996). Invasive and metastatic cells require proteases to lyse basement membrane and capillary and lymphatic walls to enter in general circulation. Previous studies have shown that a variety of proteases (such as cathepsin B, plasminogen activator, metalloproteases) are required for cancer cells to reach prostatic stroma as invasive cells and to reach distant metastatic sites (Sloane et al., 1986) (Werb, 1989) (Fidler, 1989) (Tarin, 1996) (Stetler-Stevenson and Yu, 2001). Proteases come from invasive cells, or stromal cells, or both. Morphologically, invasive cells are stem cells are undifferentiated as reported as before (Sinha et al., 1977) (Sinha, 2019). They readily pass through acinar basement membrane and colonize prostatic stroma and proliferate leading to pathological patterns described by Gleason (Gleason, 1966) (Gleason and VACURG, 1977). Invasive cells need breach the capillary wall to enter in general circulation and must exit the capillary wall before entering a distant organ (such as pelvic bones, liver, lungs, brain) for establishing metastasis. This process also requires proteases for distant organ metastasis. Stem cells produce insufficient amount proteases to lyse capillary wall to enter general circulation and to exit for metastasis. Prostatic columnar/cuboidal cells are lineage of stem cells whereas metastatic cells have dedifferentiated columnar/cuboidal cells (Sinha, 2019) (Sinha and Wilson, 2018). This led us to conclude that passage of individual invasive cancer cell beyond prostatic stroma has many barriers for a successful metastasis (Figure 1).

**Figure 1.**
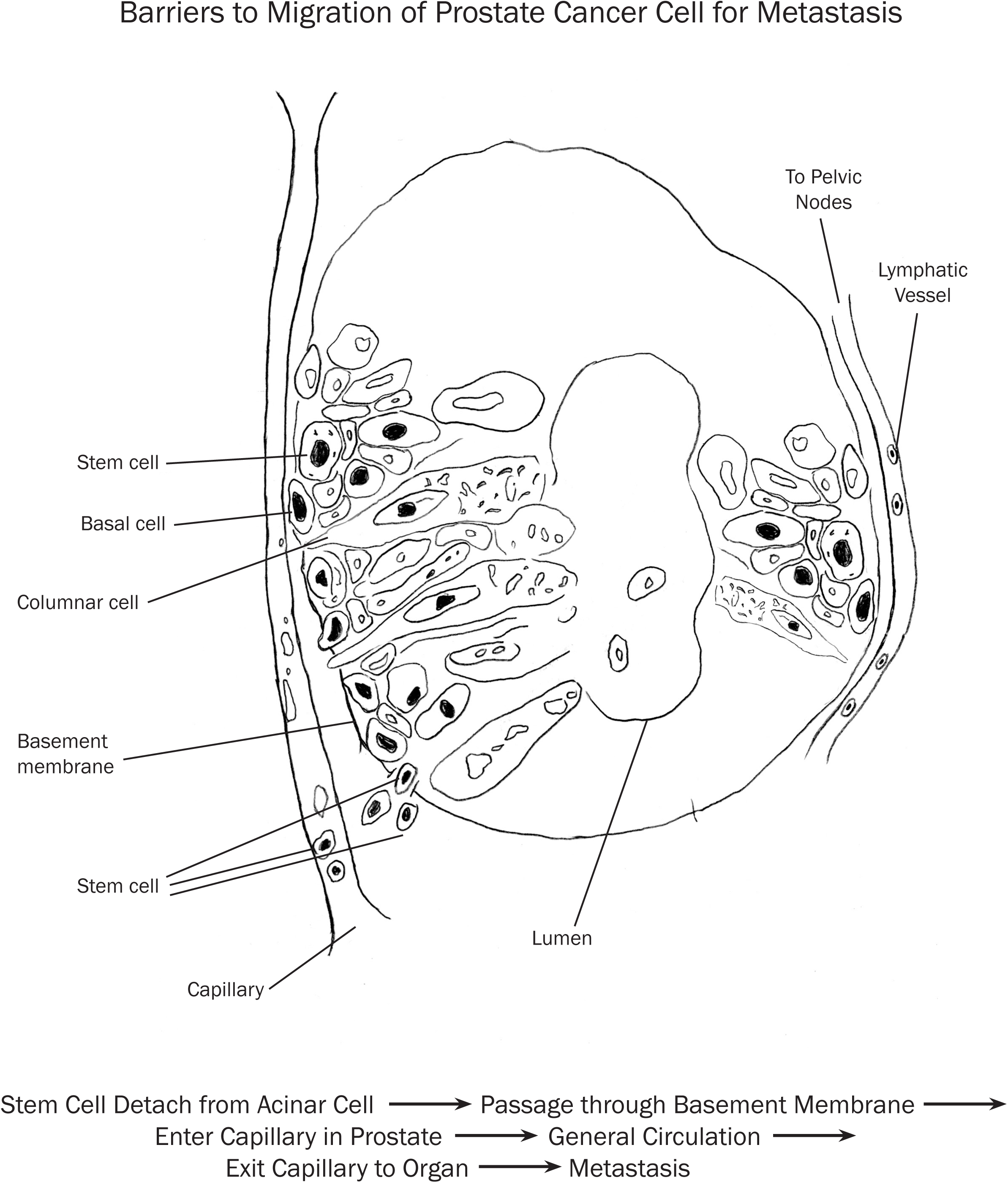
Diagrammatic figure illustrates prostatic acinus and adjacent capillary and lymphatic vessels. Diagram shows barriers to metastasis of prostate cancer cell. Each barrier requires protease (s) to lyse the membrane. See text for details.

Recent studies of Wyatt et al. showed the presence of circulating DNA which was matched in this study of tissue biopsy in prostate cancer. They suggested that DNA can be used as biomarkers (Wyatt et al., 2017). Recently, Weidle et al. showed the individual functional role of prostate cancer metastasis-related micro-RNAs (Weidle et al., 2019). This led us to hypothesize that chromatin harboring mutated DNA/genes in the nucleus of dedifferentiated columnar/cuboidal cells can readily pass through many barriers to establish metastasis whereas passage of individual cancer cell beyond prostatic stroma has many barriers to reach metastatic site (s). We have tested our hypothesis using untreated and DES (diethylstilbestrol)-treated in PC as observed by transmission electron microscopy.

## Materials and methods

Former VAMC urology surgeon, Dr. Clyde E. Blackard and his associates selected patients for biopsy and/or radical prostatectomy. Patients were not treated with any hormone therapy or chemotherapy prior to biopsy and prostatectomy. Prostate specimens were submitted to the Pathology Service of Minneapolis VAMC and specimens not used in diagnosis were collected for research between 1972 and 1975. Tissue samples were embedded in Epon 812 and stored in our laboratory. Prostate samples were obtained following the approval of the institutional review board (IRB) guidelines in place at the VA and the University of Minnesota. No University of Minnesota specimens were used in this study. We received 13 untreated samples, four BPH (benign prostatic hyperplasia) and eight DES alone or DES plus Provera treated specimens. Prostatectomy and/or biopsy tissue pieces were fixed for two hours in combination of 2% paraformaldehyde and/or 3% glutaraldehyde in 0.1M phosphate buffer at pH 7.3. Prostate pieces were washed in buffer and post-fixed in 1% to 2% buffered osmium-tetroxide, washed again, dehydrated in graded ethanol, and embedded in Epon 812 as previously described before (Sinha and Blackard, 1973) (Sinha et al., 1977). Blocks were trimmed for thick and thin sections using a Reichert-Jung microtome. Thin sections (about 400-500 angstrom) were mounted on copper grids, stained with a combination of lead citrate and uranyl acetate and examined with RCA EMU 3 or 4 electron microscopes, as detailed (Sinha and Blackard, 1973) (Sinha et al., 1977) (Sinha et al., 1987). Reynolds had shown that lead citrate was an electron-opaque stain (Reynods, 1961). Clinical details of untreated and DES-treated patients were previously published (Sinha et al., 1977). Range of DES treated cases varied from 37 days to 18 years and 9 days (Sinha et al., 1977) (Sinha et al., 2016). The age of untreated patients ranged from 58 to 79 years with a mean ± standard error of the mean of 70.54 ±3.60. The age of DES-treated patients ranged from 53 to 86 years, with a mean ± standard error of 69.37±2.83 years. Sections were graded by Drs. Donald F. Gleason and Nancy A. Staley, former staff pathologists at the Minneapolis VAMC. Patients had PC with pathological grades III and IV tumors which are comparable to Gleason histological scores 6 to 10 (Gleason, 1966) (Gleason and VACURG, 1977). Clinical stages were B, C and D (Ellis and Lange, 1994).

## Results

The prostatic stem cell has a rounded nucleus, prominent nucleolus, intact nuclear membrane, few ribosomes and small mitochondria (Figure 2a). Columnar/cuboidal cell is a lineage of stem cells and has elongated and pleomorphic nucleus in cancer cells, but not in benign prostate cells (Sinha, 2019) (Sinha and Wilson, 2018). Cuboidal/columnar cells have secretory granules, mitochondria and a portion of acinar lumen in oblique section (Figure 2a). Secretory cells are differentiated cells whereas stem cells are poorly-differentiated and have relatively few cytoplasmic organelles (Figure 2a). Inner nuclear membranes of some columnar/cuboidal cells provide a platform for anchoring intermediate filaments (Figure 2c). The inner nuclear membrane also provides areas for binding proteins for chromatin/DNA. The intermediate filaments play a role in organization of chromatin and heterochromatin and gene expression (Dittmer and Misteli, 2011) (Saarinen et al., 2015). In contrast to the nuclei of benign prostate and BPH cells, nuclei of some cancer cells loose shape or develop plasticity (or become pleomorphic) (Figure 2b). The loss of lamins and intermediate filaments results in nuclear plasticity in some columnar/cuboidal (Figure 2b). The nucleus at the top of the micrograph shows that heterochromatin is associated with the nuclear membrane and chromatin is inside the nucleus. Another nucleus shows plasticity at one end by illustrating folds in the nuclear membrane whereas the other end of this nucleus is relatively smooth. This nucleus has a prominent nucleolus. A portion of another nucleus shows folds in nuclear membranes. The nucleus at the bottom of the micrograph is completely pleomorphic and shows numerous folds and condensed nuclear material (Figure 2b). Taken together, these four nuclei illustrate the development of progressive nuclear plasticity in this PC case. Micrograph also illustrates a few nuclear folds, secretory granules and vacuoles and mitochondria whereas the other portion of the micrograph illustrates that nuclear membranes are totally pleomorphic and the boundary between nuclear membrane and cytoplasm is lost (Figure 2c). This releases nuclear material from the confines of the nuclear membrane to cytoplasm (Figure 2c). A portion of a nuclear membrane with its attached intermediate filaments is shown (Figure 2c). The nucleus shows condensed heterochromatin and chromatin (see Figure 2c). The organized structure of the nucleus is lost whereas cytoplasm still shows secretory granules and vacuoles (see Figure 2c). This brings chromatin/DNA and cytoplasm in a single compartment resulting in intermingling of their contents with cytoplasmic organelles (Figure 2c). Electron dense molecules of chromatin and/or heterochromatin are released into the cytoplasm (Figure 2c). Intermediate filaments are still attached to the nuclear membranes (Figure 2c). Chromatin harboring DNA/genes appear as electron dense molecules. Lead citrate stains basic proteins which bind DNA producing electron dense (opaque) molecules (Reynods, 1961) (Sinha et al., 1977). The later are illustrated in nucleus and adjoining cytoplasm (Figure 3a). Another micrograph shows a part of an invasive cell nucleus with electron dense molecules which are also distributed over collagen fibers (Figure 3b). Electron dense molecules are illustrated within and outside nucleus (Figure 3c). Some electron dense molecules are observed in stroma between capillary and acinar cell and in capillary endothelium and on red cell surface (Figure 3c). Figure (3d) illustrates capillary without electron dense molecules and intact nuclear membranes. Figures 3 a to c illustrate the sequence of passage of electron dense molecules to a capillary.

**Figure 2.**
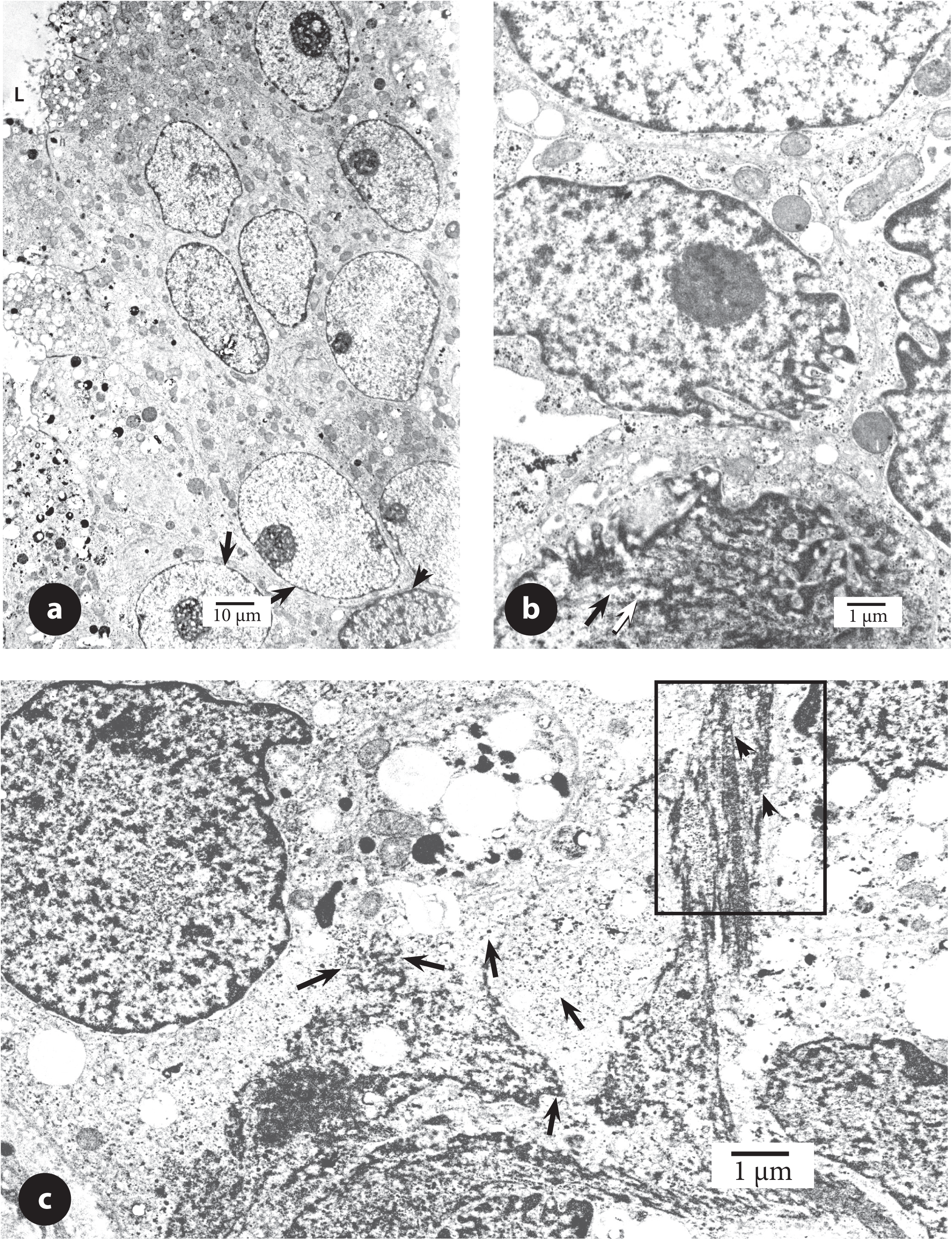
a. Micrograph shows basal cell with a spindle-shaped nucleus (arrow head) and basally located stem cells (arrows) with intact nuclear membranes, prominent nucleoli, nuclear chromatin, mitochondria, few ribosomes indicating that stem cells are undifferentiated (poorly differentiated). Oblique section shows some partially differentiated columnar/cuboidal cells with secretory granules and acinar lumen (L). Nuclei are oval to elongated, but not pleomorphic in acinar cells. (untreated patient # 110). Bar shows magnification. b. Figure illustrates four nuclei of columnar/cuboidal cells. mitochondria, ribosomes and some secretory granules, all of them are usually found in dedifferentiated cells. The nucleus at the top of the micrograph shows smooth nuclear membrane associated with heterochromatin and chromatin. Another nucleus shows plasticity in the nuclear membrane as illustrated by folds whereas the other end of this nucleus is still smooth. Portion of another nucleus shows several folds. The nucleus at the bottom of the figure is completely pleomorphic. Some chromatin electron dense molecules have been released in the cytoplasm (arrow in the boxed area). Taken together, these four nuclei illustrate development of nuclear plasticity. (untreated patient #117). The bar shows magnification. c. Figure illustrates a nucleus with intact nuclear membrane and condensed chromatin and heterochromatin and a set of three nuclei which have lost their shape and boundary between nuclear membranes and cytoplasm. Heterochromatin and chromatin appear as electron dense molecules (arrows). Portions of intermediate filaments are illustrated (arrow heads, area enclosed by a rectangle) as shown in Figure 3d. (untreated patient # 114). The bar shows magnification.

**Figure 3.**
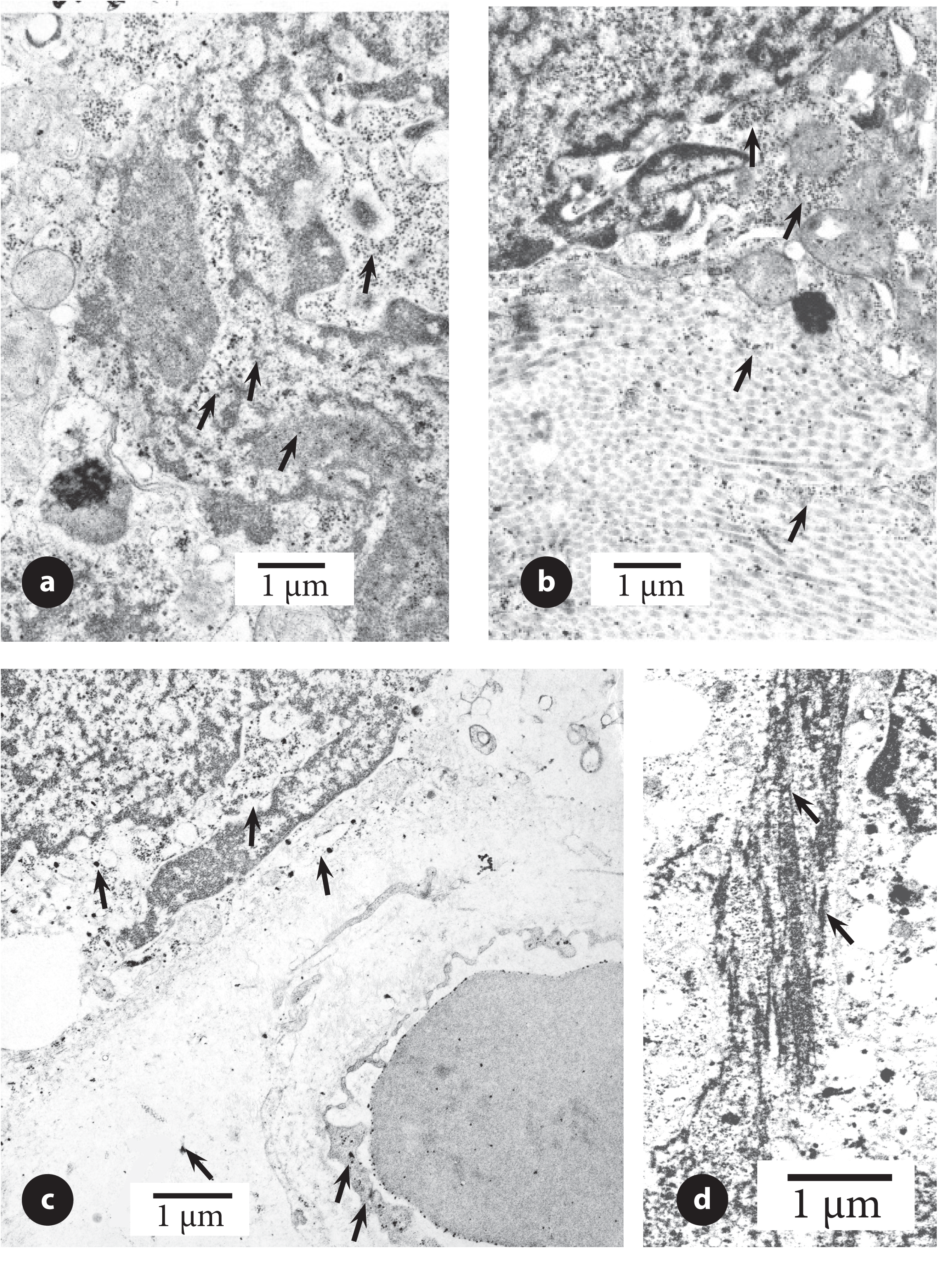
a.This micrograph illustrates an advanced stage of nuclear plasticity. Aggregates of nuclear chromatin/DNA appear as clusters of discrete electron dense molecules (arrows) in nucleus and cytoplasm. (untreated patient # 110). The bar shows magnification. b. A metastatic cell with pleomorphic nucleus in the stroma is surrounded by collagen fibers. The nucleus illustrates condensation of chromatin as electron dense molecules in cytoplasm and on collagen fibers (arrows). (untreated patient #110). The bar shows magnification. c. Pleomorphic nucleus illustrates electron dense molecules. These molecules are in stroma, in capillary, capillary endothelium (arrows) and on red cell surface. Arrows indicate that some electron dense molecules have been transported to the stroma, capillary endothelium and red cell. (untreated patient # 114). The bar shows magnification. d. Details of intermediate filaments shown in Figure (2c). The figure illustrates that electron dense molecules are associated with intermediate filaments (arrows). The bar shows approximate magnification.

The nuclear plasticity was observed in untreated and DES-treated PC, but not in benign (normal) prostate and BPH. In DES-treated cases, chromatin/DNA appeared as electron dense molecules which were released from the nucleus to cytoplasm much as in untreated cases (Figure 4d). Nucleolus was present in DES treated cases. Nuclear membranes in adjacent acinar cells did not show plasticity (Figure 4a). This capillary was not near a pleomorphic nucleus. Metastatic cell nucleus is distinctly different from dying cell (cell death). Cell death has condensed nuclear chromatin and heterochromatin and degenerated cytoplasmic organelles (Figure 4b). Adjacent acinar cells had not degenerated and have cytoplasmic organelles and nuclei comparable to those shown in Figure (2a). The loss of nuclear membrane between nucleus and cytoplasm allows release of electron dense molecules from the confines of nuclear membranes into cytoplasm then in stroma and finally in nearby circulation. These molecules are carried to the capillary as shown by a series of micrographs (Figures 3a-c, 4c-e) and presumably to metastatic sites. Once in circulation, electron dense molecules can reach and colonize several organs (such as liver, lung, pelvic bones, and/or brain). We have not studied lymphatics for the presence or absence of electron dense molecules.

**Figure 4.**
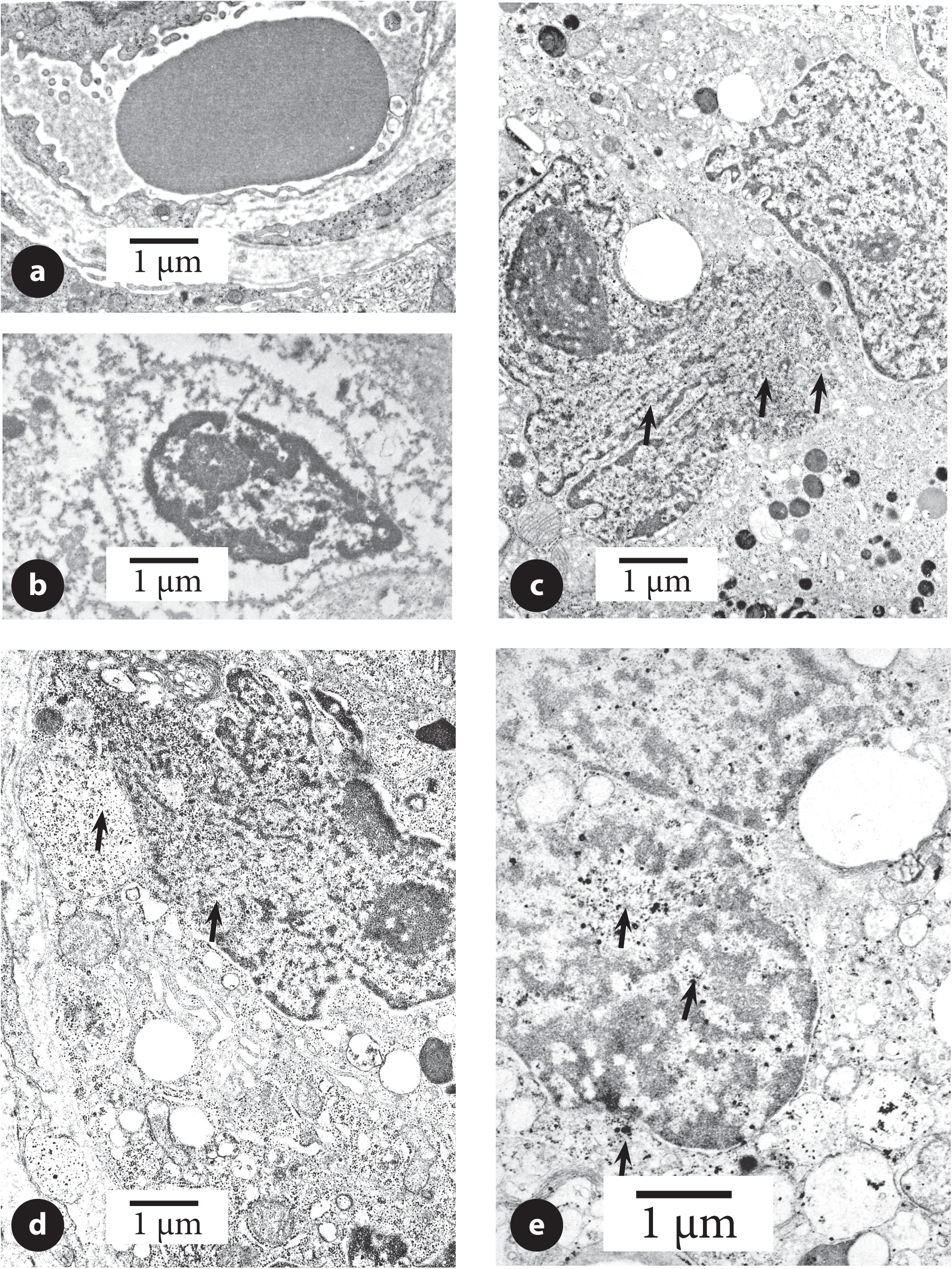
a. A capillary with red blood cell, endothelium, stromal connective tissue and a portion of acinar cells, Capillary was not near any pleomorphic nucleus and does not show electron dense molecules in stroma and capillary endothelium. DES-treated for 37 days (patient # 118). The bar shows magnification. b Figure illustrates cell death with pyknotic nucleus and cytoplasm that has lost most of its organelles. Acinar lumen contained sloughed cytoplasmic portions. Pyknotic nucleus has condensed nuclear material and thickened nuclear membranes. Nucleus still contains nucleolus. Adjacent columnar/cuboidal cells did not show any signs of degeneration in nuclei and cytoplasm. This patient was treated with DES for 37 days prior to biopsy (patient #118). The bar shows magnification. c. A light microscope figure of an acinus shows a migrating invasive cell to stroma (arrow). Another arrow indicates acinar cells in stroma. Acinar lumen has several sloughed cells in lumen. The bar shows magnification. d. In a cell, nuclear plasticity is illustrated by loss of the nuclear membranes. Some electron dense molecules (arrows) are present in the nucleus and cytoplasm. This nucleus has a large nucleolus. This patient was treated with DES for 37 days prior to biopsy (patient #118). The bar shows magnification. e. Figure illustrates portions of two nuclei with condensed chromatin and heterochromatin. Electron dense molecules in nucleus intermingle with cytoplasmic organelles. Some of the dense molecules are present in cytoplasm (arrows) and within nucleus area. This patient was treated with DES for 18 years and 9 days prior to biopsy (patient #104). The bar shows magnification.

## Discussion

Several studies have shown that DNA is shed into the blood stream of advanced metastatic castration resistant prostate cancer (CRPC) and this circulating DNA can be used as a marker (Wyatt et al., 2017) (Ritch and Wyatt, 2018) (Annala et al., 2018). CRPC is u)niformly fatal disease (Van Etten and Dehm, 2016). These studies did not identify (or categorize metastatic cell. Our analysis of electron microscopic study has shown that metastatic cell is identified by a) nuclear plasticity, b) loss of nuclear membranes, c) lack of boundary between nucleus and cytoplasm and d) electron dense molecules). All these features are found only in metastatic cells and not in stromal invasive cells. The presence of one or two features is inadequate to identify metastatic cell in tissue sections. This also led us to further investigate the most important feature that can be utilized in diagnosis of metastatic cancer in tissue section. Our review has also shown that separation of nuclear and cytoplasm compartments is critical for the functioning of cells in benign prostate and PC and other cancers (Raska et al., 1990) (Dittmer and Misteli, 2011) (Saarinen et al., 2015). The loss of lamins and intermediate filaments leads to pleomorphic (or plasticity) in nuclei of some columnar/cuboidal cells (Prokoccimer et al., 2009) (Pajerowski et al., 2017). Nuclear morphometry at light microscopic level has been related to Gleason grade and used as predictor of prognosis (Diamond et al., 1981) (Partin et al., 1989). We conclude that the lack of boundary between nucleus and cytoplasm is the single most important feature of a metastatic cell. At the present, electron microscopy is the best approach for identifying metastatic cell.

Chromatin/DNA are small molecules, much as nutrients, metabolites, viruses, bacteria, drugs; all of them readily cross many barriers. Small molecules readily move-in and out of cells unlike an individual prostate cancer cell which faces several barriers (Figure 1). We have not shown the presence of chromatin /DNA (electron dense molecules) at metastatic sites. but have provided morphological evidence that molecules reach the capillary and red cell surface. In conclusion, PC has at least two sub-populations of cells with and without nuclear plasticity. Since our study is based upon small number of samples, it needs to be confirmed by others. Our analysis of metastatic process in the prostate contrasts with the numerous previous studies showing that migration of individual cancer cell (s) produce metastasis (Stetler-Stevenson and Yu, 2001) (Killion and Fidler, 1989) (Heppner and Miller, 1989) (Liotta and Stetler-Stevenson, 1991).

The castration resistant prostate cancer patients indicate a failure of androgen deprivation therapy (ADT) and of Abiraterone and/or Enzalutamide treatments (Wyatt et al., 2017) (Ritch and Wyatt, 2018) (Annala et al., 2018) (Tucci et al., 2019) Recently, Crawford et al. have traced the progress made by a variety of antiandrogen treatment and have advocated its continued use in the treatment of PC (Crawford et al., 2018). Earlier treatments of PC by hypophysectomy and/or adrenalectomy were performed in patients who no longer responded to ADT or similar treatments (Huggins and Scott, 1945) (Brendler, 1973) (Walsh, 1975) (Silverburg, 1977). The current and previous efforts provided limited benefits to patients who ultimately succumbed to PC. These treatments were usually administered to patients on the assumption that some residual androgen and androgen-dependent cells were still present. These studies indicate that efforts to treat PC with ADT treatment has failed for over fifty years. Androgen dependence of prostate has a long history and has many advocates who do not support change despite several studies showing the presence of estrogen localization in PC (Sinha et al., 1973) and androgen, estrogen and progesterone receptors in prostate and breast cancers (Bashirelahi et al., 1983) (Greene et al., 1984) (Hu et al., 2012). In recent years, we have shown estrogen receptors emerged in patients who had a failure of ADT treatment (Sinha et al., 2016). We suggest that CRPC should also be treated with tamoxifen, aromatase inhibitors and/or similar drugs.

### Future direction of metastasis research and treatment

The selection of metastatic site (s) is a random and/or semi random process. For example, PC usually metastasizes to pelvic bones, liver, lungs, and brain. We postulate that mutated nuclear DNA of PC enter the nuclei of host cells and induce host cells to produce metastatic PC. Mutated genes have proliferation advantage whereas non-mutated genes do not. Presence of mutated prostate genes, especially in aggressive CRPC, in host cells (such as liver) can also induce some liver cell genes to proliferate resulting in liver cancer. We have not shown in this study, but it would suggest the presence of metastatic PC in liver and liver cancer in liver. While metastatic PC is treated, liver cancer remains untreated. Both types of cancers need to be treated for a successful outcome of metastatic disease. Whether two types of cancers are mixed together or remain separated are also unknown. A similar scenario probably exists for metastatic PC in lungs, pelvic bones and/or brain. A similar case can be made for breast, lung and other solid organ cancers and each cancer needs to be explored separately. Our study provides some of the reasons for the failure of treatment for metastatic PC. This also explains why the efforts of so many scientists and clinicians have failed for so many decades to successfully treat metastatic cancers. Our idea can be readily assessed by using concurrent localization of markers for prostate and liver cancers. Our idea also needs to be explored further.

## Conflicts of Interest

The Author has no conflict of interest to report regarding the publication of this article.

## Disclaimer

The opinion expressed in this article is that of the Author and not of the U.S. Government, Department of Veterans Affairs or the University of Minnesota.

## Acknowledgements

This research was supported in part by the Research Service of the Minneapolis Veterans Affairs Medical Center by providing laboratory and other research facilities to AAS. We are grateful to Dr. Donald F. Gleason and Dr. Nancy A. Staley, former pathologists of the Minneapolis VA Medical Center, for grading prostate cancer sections. We are also grateful to Dr. Clyde E. Blackard and his associates for biopsy and prostatectomy specimens. The Author thanks Mr. Francis F. Pomroy, Jr. formerly of the Minneapolis VA Medical Center for making sections for this study. The author is grateful to James S. Hungaski, Jr. and Mr. Jonathan Erickson of the VAMC Media Service for making the final figures, and to the staff of the Departments of Surgical Pathology, Library, and the Research Service, Minneapolis VA Medical Center. The Author thanks Ms. Martha K. Grace for helpful comments and critical proof reading of the manuscript.

